# Direct cerebellar control over motor production in a species with extreme cerebellar enlargement

**DOI:** 10.1101/2025.03.10.642493

**Authors:** Federico Pedraja, Michael Genecin, Dillon Noone, Dan Biderman, Philip Cho, David E. Ehrlich, Nathaniel B. Sawtell

## Abstract

The cerebellum is thought to fine-tune movement without being required for its production. However, this textbook view derives mainly from studies of mammalian species with highly developed cerebral cortices. Here we examined cerebellar function in the elephant-nose fish, a member of a family of African weakly electric fish (*Mormyridae*) in which the cerebellum is massively enlarged. The elephant-nose fish is named for a flexible facial appendage that is used to probe surfaces and extract prey from substrate. Results from microstimulation, electrophysiological recordings, and lesions support a direct role for the C1 region of the mormyrid cerebellum in controlling movement of this appendage. These findings suggest that the cerebellum is capable of performing functions typically ascribed to the cerebral cortex, emphasizing the importance of evolutionary history on the functional specialization of brain regions.

## Introduction

The brain of mormyrid fish constitutes roughly 3% of their body weight (comparable to 2-2.5% for human brains) and accounts for an estimated 60% of resting oxygen consumption (compared to 20% in humans and 2-8% in most other vertebrates) ^1-4^. This extreme encephalization is primarily due to the expansion of the cerebellum, which covers the entire dorsal surface of the brain in many mormyrid species ^5,6^ (**Fig. 1A**). Anatomical and *in vitro* electrophysiological studies have shown that this “gigantocerebellum” is similar to the cerebellum of other vertebrates in terms of its cell types, microcircuitry, gene expression patterns, and basic electrophysiological properties ^7-12^. For example, like their mammalian counterparts, mormyrid Purkinje cells integrate tens of thousands of excitatory parallel fiber inputs onto their spiny apical dendrites with a single, powerful excitatory input from the inferior olive. However, despite its long-held fascination, the function of the mormyrid cerebellum remains virtually unexplored.

**Figure 1.**
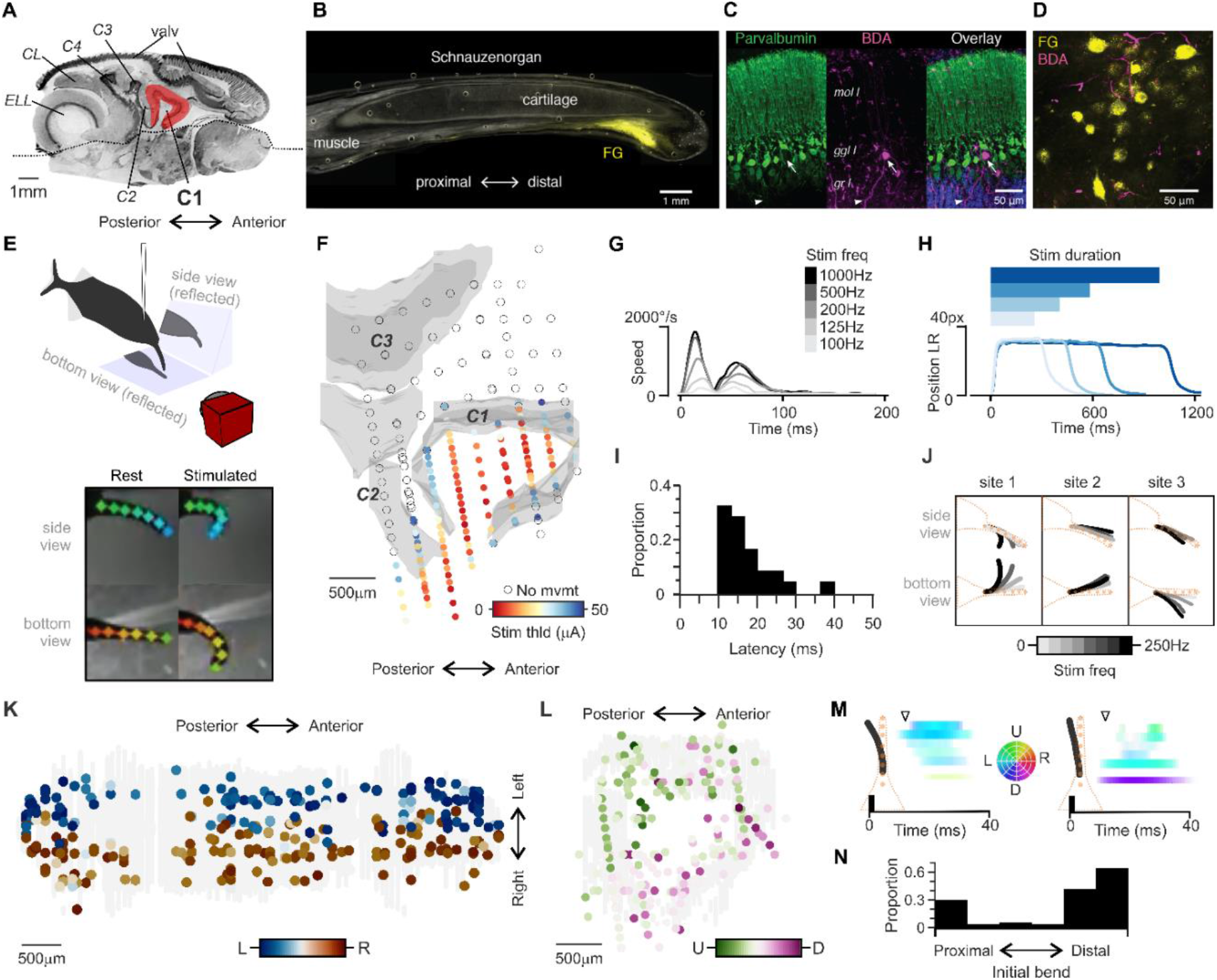
C1 microstimulation evokes SO movements. **(A)** Parasagittal section of the brain of *Gnathonemus petersii*. All structures above the dotted line are cerebellum or cerebellum-like. Abbreviations: electrosensory lobe (ELL), caudal lobe (CL), valvula (VA), C (corpus). **(B)** Cross-section of the SO following an intramuscular injection of the retrograde tracer fluorogold (FG). **(C)** Transverse section through C1 showing Purkinje cells stained with an antibody to parvalbumin (green) and C1 efferent neurons labeled by a biotinylated dextran amine (BDA) injection into C1 (magenta). Selective BDA labeling of efferent neurons in this case is likely due to injection site placement. **(D)** C1 axon terminals (BDA, magenta) are found in the vicinity of FG-labeled trigeminal motor neurons after dual tracer injections in the SO and C1. **(E)** Setup for C1 microstimulation and multi-view SO tracking in head-fixed fish. **(F)** Reconstruction of microstimulation sites from a single fish. The color code indicates the threshold stimulus intensity for evoking an SO movement. Empty circles indicate threshold >50 µA. **(G)** Effects of grading stimulation frequency on SO movement speed at a single site. **(H)** Effects of grading stimulation duration on SO position at a single site. **(I)** Minimum latency to SO movement onset (n = 22 sites). **(J)** Examples of site-specificity of SO movements evoked by graded stimulation. **(K)** Spatial organization of leftward versus rightward movement components. The Purkinje cell layer of C1 is represented as a flattened sheet (n = 328 stimulation sites from 5 fish). **(L)** Spatial organization of upward versus downward movement components. Sites are mapped onto a single mid-sagittal section through C1 (n = 328 stimulation sites from 5 fish). Only sites within 200 µm of the Purkinje cell layer are included in **K**,**L. (M)** Effects of SO stimulation over time at two sites illustrating movement initiation at distal (left) versus proximal locations along the SO. Arrowhead indicates the time illustrated in the schematics and in **N. (N)** Histogram showing the proximo-distal location of SO movement initiation (n = 800 stimulation sites from 4 fish).

## Results

To address this, we examined the C1 region of the mormyrid cerebellum in the species *Gnathonemus petersii* (commonly known as the elephant-nose fish). The C1 region is particularly intriguing because it makes a prominent excitatory projection to the motor nucleus that controls the finger-like facial appendage, or schnauzenorgan (SO) ^9,13^. The SO is a flexible cartilaginous cylinder protruding from the dental bone, encased by longitudinally oriented muscles with attachments spanning its entire length ^14,15^. Activation of muscles at different radial locations control movement direction in 360 degrees whereas activation of muscles at different proximal-distal locations allows for finely articulated, finger-like bending. The SO is densely covered by electroreceptors and massively innervated by the sensory trigeminal nerve and functions both as a mobile sensor used to identify and localize invertebrate prey in murky rivers and an actuator capable of finely articulated movements used for extracting them from the substrate ^14-20^. We performed additional anatomical tracing experiments by injecting a fluorescent retrograde tracer in SO muscles and a second anterograde axonal tracer in C1 (**Fig. 1B,C**). Note that glutamatergic cerebellar output neurons in teleost fish (including mormyrids) are intermingled with Purkinje cells rather than being located in a separate deep nucleus as in most other vertebrates ^2,21^. Labeled axons were observed in close proximity to SO motor neurons in the ipsilateral motor trigeminal nucleus (**Fig. 1D**), consistent with C1 projections directly targeting motor neurons that control SO movement.

### C1 microstimulation evokes SO movements

To make a functional link between C1 and SO movement we performed electrical microstimulation in head-restrained fish combined with multi-view high-speed videography to track SO movements (**Fig. 1E**). Brief trains of low-intensity stimulation elicited SO movements in C1, but not neighboring cerebellar regions (**Fig. 1F**). Near threshold stimulation at anterior sites in C1 typically elicited SO movement only, while more posterior sites sometime also evoked tail and fin movements (**Fig. S1**). The amplitude and speed of SO movements exhibited a consistent relationship with microstimulation parameters independent of the location of the stimulation site within C1. SO movement amplitude and speed increased monotonically with stimulation current strength and pulse train frequency (**Fig. 1G, Fig. S1**). SO movement amplitude and duration also increased monotonically as pulse train duration was increased (**Fig. 1H**). These characteristics, together with the short latency of the evoked movements (**Fig. 1I**), further support direct effects of C1 output on SO motor neurons.

Movement direction (quantified from the maximum horizontal and vertical displacement of the tip of the SO) varied systematically depending on the location of the stimulation site within C1 (**Fig. 1J**). The horizontal component of SO movements was typically ipsiversive to the stimulation site, i.e. stimulation on the left side of C1 tended to evoke leftward movements (**Fig. 1K**). This is consistent with ipsilateral projections from C1 to the trigeminal motor nucleus and from motor neurons to SO muscles ^9,13,14^. A tendency was also observed for dorsal stimulation sites to evoke movements with relatively larger upward components and ventral sites to evoke movements with relatively larger downward components (**Fig. 1L**). Finally, stimulation at different sites produced movements that initiated at different proximo-distal points along the SO, mainly at extreme proximal or distal locations (**Fig. 1M-N**). No obvious topographical organization was observed in relation to this proximo-distal axis. Overall, these results strongly support a direct role for C1 in fine-grained control over SO movement.

### C1 neurons encode SO movement-related information

Prior *in vivo* recording from the mormyrid cerebellum have focused on characterizing electrosensory responses in the caudal lobe and portions of the valvula ^22,23^. To provide an initial description of the response properties of C1 neurons, we performed Neuropixel probe (*NP Ultra*) recordings in C1 of awake, head-fixed fish ^24^ (**Fig. 2A**). C1 neural activity was measured in response to a battery of stimuli including passive displacements of the SO, passive displacements of the tail, electrosensory stimulation of both the SO and non-SO skin regions, and electric organ corollary discharge (EOCD) input related to EOD motor commands emitted spontaneously by the fish. Passive SO displacements were by far the most effective stimulus (**Fig. 2B**), consistent with anatomical evidence for a prominent pathway from myelinated sensory trigeminal nerve fibers innervating SO muscle tendons to C1 and with electrophysiological recordings from trigeminal nerve fibers demonstrating responses to SO position and velocity ^9,15^. The paucity of electrosensory and EOCD responses we observed may reflect the lack of direct input to C1 from regions associated with electrosensory processing ^9^ and/or a requirement for more realistic electrosensory stimuli.

**Figure 2.**
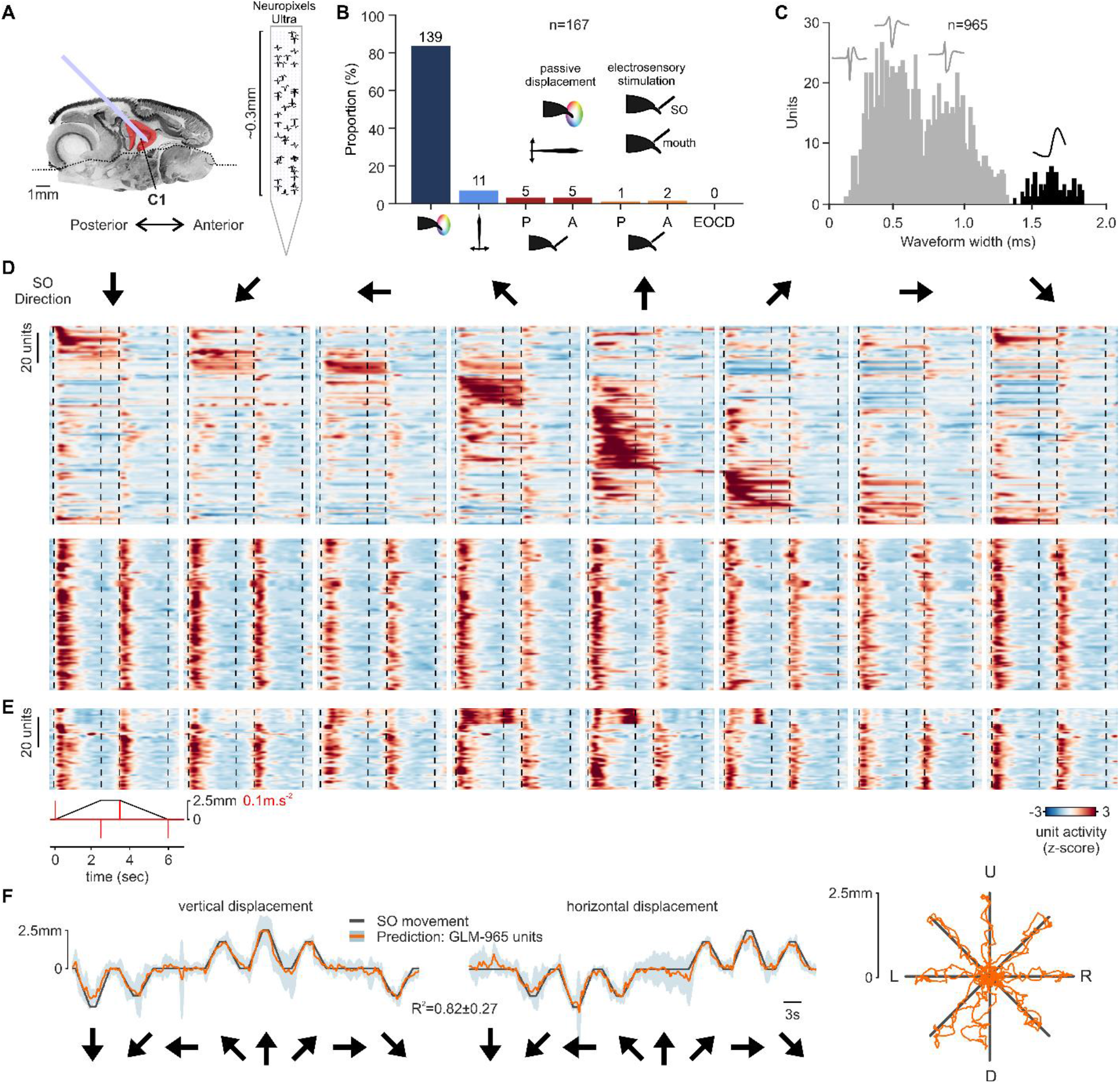
*In vivo* electrophysiological characterization of C1. (**A**) Left, schematic of Neuropixels Ultra recording in the C1 of an immobilized fish. Right, example waveforms from 48 simultaneously recorded single-units in C1 shown at their measured location on the probe. (**B**) Proportion of units responding to: passive displacement of the SO in eight directions using a motorized micromanipulator, passive lateral displacement of the tail, local electrosensory stimulation of the SO or mouth to activate either the passive (P) or active (A) electrosensory systems, and electric organ corollary discharge input related to the motor command to discharge the electric organ (EOCD). (**C**) Distribution of spike waveforms for units responsive to SO displacement (n = 965 units from 4 fish). Broad-spiking units (black) form a distinct class likely corresponding to complex spikes of Purkinje cells. (**D**) Z-scored firing rate responses of narrow-spiking units to SO displacement (each column represents a different movement direction indicted by arrows). Clustering of responses revealed units that exhibited tuning to movement direction (top, n = 141 units) and others that appeared to encode acceleration (bottom, n = 571). Units are ordered by their time of peak activity. Dashed black lines correspond to movement acceleration and deceleration. (**E**) Same as (**D**), but for all recorded broad-spiking units (n = 45). (**F**) Vertical (left) and horizontal (middle) SO position trajectories along with the mean and standard deviation of generalized linear model (GLM) fits (orange) based on the spiking of 965 recorded units from 4 fish. Right, 2D representation of both vertical and horizontal movement directions (black) along with the GLM fits.

For further analysis of responses to SO displacement we sub-divided C1 units into two classes based on spike waveform characteristics (**Fig. 2C**). A distinct class of units exhibited broad spike waveforms and low firing rates (broad: 2.7 ± 1.5 Hz; narrow: 10.5 ± 16.2 Hz). These units likely corresponded to Purkinje cell complex spikes which are commonly recorded in the molecular layer ^25^. Prior *in vitro* and *in vivo* studies using intracellular recordings have shown that mormyrid Purkinje cells share basic characteristics with Purkinje cells in other vertebrate cerebella: firing both simple and complex spikes, the latter due to a powerful excitatory synaptic input from the inferior olive ^22,26^. The remaining majority of recorded units could not be definitively sub-classified based on their spike waveforms and auto-correlograms and are simply referred to as narrow spiking units.

Diverse responses to SO displacement were observed across C1 units (n = 965), including a subset of narrow spiking units that exhibited clear directional tuning evident during the ramp and hold phase of the displacement (**Fig. 2D**). Another common response type observed in both narrow and broad spiking units encoded acceleration (**Fig. 2D,E**). A clear spatial mapping of response types within C1 was not observed. For example, units that responded most strongly to leftward movements were found on both sides of C1. The lack of such a relationship could be due to differences in how SO sensors are engaged by passive displacements versus active movements. Alternatively, mossy fiber inputs from trigeminal sensory nuclei may form widely divergent connections within C1, such that a diverse array of SO movement and position information is available for sculpting the output of each location on the C1 map. Consistent with the latter possibility, mossy fibers are a frequently encountered unit type in Neuropixel recordings from mouse and primate cerebellum ^25^.

We used a generalized linear model (GLM) to fit SO displacement trajectories using C1 neural responses. SO trajectories could be accurately reconstructed based on the recorded responses (**Fig. 2F**), with a major contribution from the population of direction selective narrow-spiking units (R^2^=0.77±0.12; n = 141) and a smaller contribution from acceleration tuned units (R^2^=0.49±0.19; n = 571). Many motor control structures, including primary motor cortex, receive prominent proprioceptive feedback from the muscles they control ^27^. Hence the observation that C1 contains rich representations of SO movement based on sensory feedback further supports a role for C1 in controlling SO movement.

### C1 lesions result in hemi-paresis of the SO

Finally, we sought to directly test the necessity of C1 for SO movement using lesions. We constructed a chamber inside the fish’s home tank enabling us to obtain high-speed video recordings of foraging behavior using three camera views. After a habituation period, fish voluntarily entered the chamber to forage for prey (live black worms) placed singly either directly on the chamber floor or inside a narrow rectangular trough (**Fig. 3A**). Fish used their SO to locate prey in both conditions and to extract them from the trough in the latter condition. Machine learning algorithms were used to reconstruct the pose of the fish in three dimensions (including 5 points along the length of the SO) (**Fig. 3B, Video S1**). Foraging behavior was recorded for at least 5 sessions before and after lesioning C1. Partial lesions of C1 were performed by passing current through a metal microelectrode positioned at sites within C1 at which SO movements could be evoked with low stimulation currents and subsequently verified histologically (**Fig. 3C**). Although a decrease in movement speed was observed following C1 lesions (**Fig. S2**), basic swimming patterns along with the motivation to forage and the ability to detect prey appeared normal, as indicated by comparable rates of worm intake in the floor condition following C1 lesions (**Fig. 3D**). In contrast, the ability of fish to extract worms from the trough exhibited a persistent post-lesion decrease (**Fig. 3E**). Kinematic trajectories revealed a persistent post-lesion decrease in the curvature and range of motion of the SO, which was particularly evident in the yaw axis (**Fig. 3F-G,I-J**). The most severely impacted animals exhibited a nearly complete paralysis of the SO (**Fig. 3F,G**) (**Videos S2, S3**). Fish with predominantly unilateral lesions, exhibited a contralateral shift in the resting position of the SO (**Fig. 3H**), consistent with a loss of muscle tone due to removal of C1 excitatory drive to SO motor neurons. The magnitude of such effects is likely underestimated by our analysis, since we were unable to distinguish between active movements of the SO versus passive bending due to frequent contact with the floor or trough during foraging. In contrast to the clear effects on the SO, the range of motion of other body parts, including the tail, were relatively unaffected (**Fig. 3K-L**).

**Figure 3.**
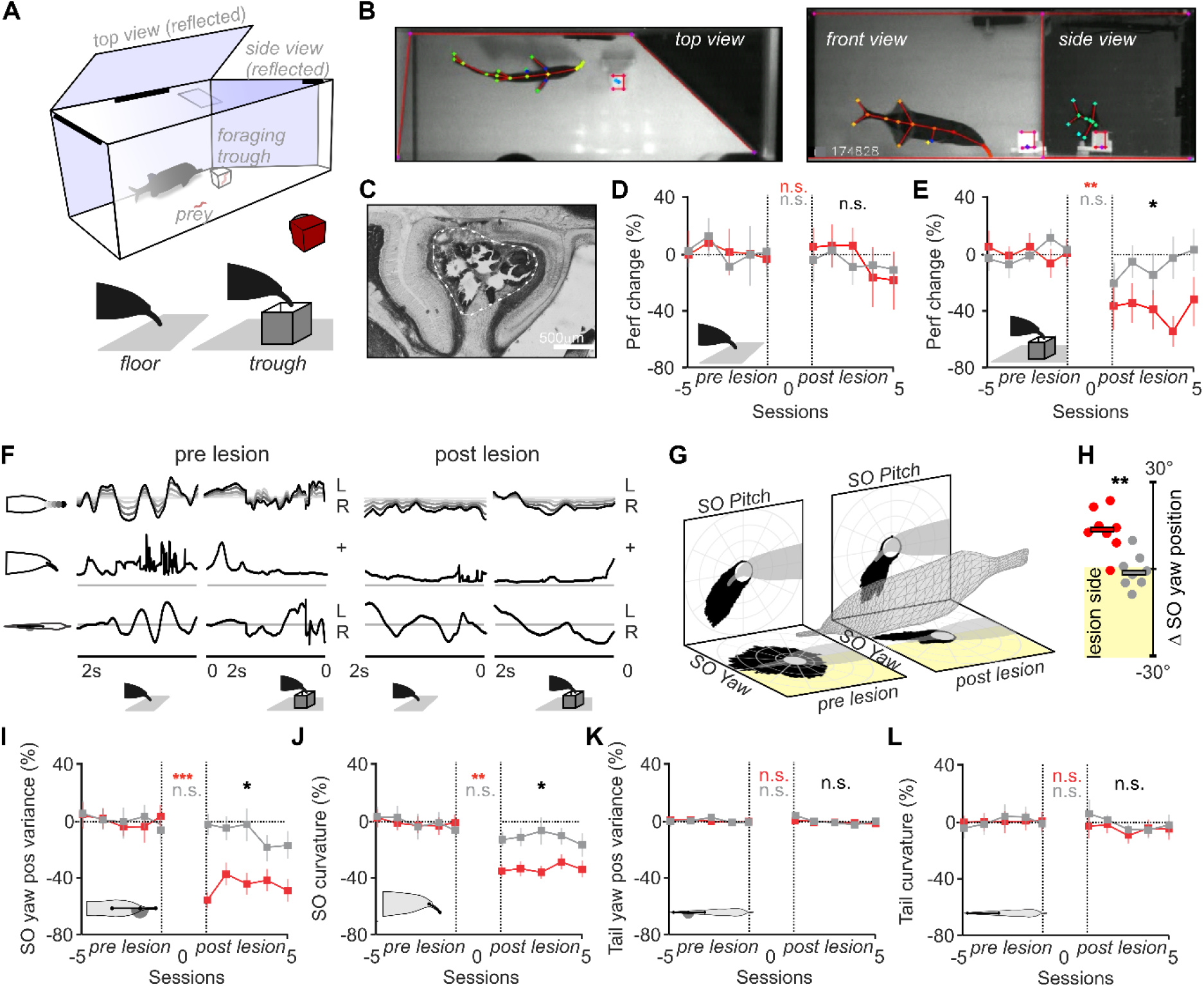
Effects of C1 lesions on foraging behavior and SO movement. (**A**) Experimental setup for multi-view recording of foraging behavior. (**B**) Three camera views with labeled keypoints used for 3D pose estimation. (**C**) Parasagittal section from a C1 lesioned fish. White dashed line marks the lesioned area. (**D-E**) Foraging performance (successful captures/total attempts) under floor (**D**) and trough (**E**) conditions for five consecutive before and after C1 (red, n = 9) or sham lesion sugery (gray, n = 8). Note the decline in foraging performance after C1 lesion in the trough condition (Floor: post C1 lesion vs post sham-lesion: Mann-Whitney test, p=0.64; pre vs post C1 lesion: Wilcoxon test, p=0.38; pre vs post sham lesion: Wilcoxon test, p=0.56. Trough: post C1 lesion vs post sham-lesion: Mann-Whitney test, p=0.03; pre vs post C1 lesion: Wilcoxon test, p=0.003; pre vs post sham lesion: Wilcoxon test, p=0.85). (**F**) Kinematic traces showing SO position for different proximo-distal segments (grayscale) (top), SO curvature (middle), and tail position (bottom) during foraging under floor and trough conditions for an example fish before and after C1 lesion. (**G**) Distribution of SO pitch and yaw angle distribution before (left) and after C1 lesion (right). Same fish as in (**F**). Yellow shading indicates the lesion side. (**H**) Change in average SO position (pre vs. post-lesion) for fish with lateralized C1 lesions (n = 8) and sham-lesion controls. Mann-Whitney test: p=0.002. (**I-L**) SO yaw position variance, curvature, and tail yaw position variance and curvature before and after C1 (n = 10) or sham (n = 8) lesions. Range of motion of the SO decreased after C1 lesions while that of the tail was unchanged (SO yaw position variance: post C1 lesion vs post sham-lesion: Mann-Whitney test, p=0.02; pre vs post C1 lesion: Wilcoxon test, p=0.001; pre vs post sham lesion: Wilcoxon test, p=0.31. SO curvature: post C1 lesion vs post sham-lesion: Mann-Whitney test, p=0.04; pre vs post C1 lesion: Wilcoxon test, p=0.005; pre vs post sham lesion: Wilcoxon test, p=0.78. Tail yaw position variance: post C1 lesion vs post sham-lesion: Mann-Whitney test, p=0.39; pre vs post C1 lesion: Wilcoxon test, p=0.89; pre vs post sham lesion: Wilcoxon test, p=1.00. Tail curvature: post C1 lesion vs post sham- lesion: Mann-Whitney, p=0.84; pre vs post C1 lesion: Wilcoxon test, p=0.70; pre vs post sham lesion: Wilcoxon, p=0.64).

## Discussion

The mormyrid cerebellum has been well-known to comparative neuroanatomists for over a century due to its size and histological regularity ^28^, yet its function remains mysterious. While there is clear evidence linking some parts of the mormyrid cerebellum to electrosensory processing ^7,23^, there are also large regions (including C1) that do not receive electrosensory input. Furthermore, South American weakly electric fish possess highly developed electrosensory and electromotor abilities, similar to those of mormyrids ^29^, but show much less cerebellar enlargement. The present study provides evidence from microstimulation, electrophysiological recordings, and lesions that the C1 region of the cerebellum in *Gnathonemus petersii* plays a critical role in controlling the SO, most likely mediated by direct excitatory projections to trigeminal motor neurons innervating SO muscles. A direct role for the cerebellum in motor production is intriguing because it contrasts with the textbook view that motor cortex is primarily responsible for motor command generation while the cerebellum performs fine-tuning or online correction based on errors ^30-32^. These distinct functional roles for cerebral versus cerebellar circuitry have typically been assumed to reflect fundamentally different computational capacities of the “deep”, highly recurrent circuitry of the former as compared to the “shallow”, feedforward circuitry of the latter ^33^.

An alternative suggested by the present study is that the core computational capacities of cerebral and cerebellar cortical circuitry are highly overlapping, with evolutionary history playing a decisive role in their functional specialization. According to this view, cerebral functions would be expanded in species in which natural selection favored physical enlargement of the cerebrum (e.g. primates), whereas cerebellar function would be expanded in a species such as *Gnathonemus petersii*, in which the cerebellum is enlarged relative to other brain structures ^5,6^. Consistent with this, partial paralysis of the SO observed after C1 lesions appears more similar to the effects of motor cortex lesions on forelimb use in primates than to conventional cerebellar ataxia ^34,35^. While an obligatory role for the cerebellum in the initiation of primate smooth pursuit eye movements is well-established ^36^, other types of eye movement, including saccades can still be generated ^37^. In the case of C1 lesions, all SO movement, including “spontaneous” SO movements generated when the fish is far from prey, appear to be strongly impacted. Additional studies of the mormyrid cerebellum, including C1 recordings in freely swimming fish ^38^, provide an ideal test bed to understand whether and how feedforward cerebellar architecture supports continuous real-time control over a relatively high degree of freedom end-effector. A question of particular interest is how prominent sensory feedback to C1 is leveraged for control.

While the present study focuses exclusively on C1, other brain regions are likely to be important in SO control and foraging behavior more broadly. The trigeminal motor nucleus receives direct input from a number of structures in addition to C1, including a region of the distal valvula (the cerebellar sub-region that shows the greatest enlargement in mormyrids) and the optic tectum (homologous to the superior colliculus) ^13^. Extensive studies in mammals suggest that the superior colliculus and the cerebellum work in concert to control sensory guided movements, e.g. saccadic eye movements ^39^. Interestingly, direct projections from the forebrain to the trigeminal motor nucleus (i.e. the analog of a descending motor cortical pathway), have not been reported ^13^. Finally, it is intriguing from an evolutionary and comparative perspective that mormyrids exhibit remarkable diversity in craniofacial morphology ^40^. In addition to the mobile SO, some species have elongated frontal bones with small pincer-like jaws at the anterior tip of the rostrum (known as tubesnouts), others have tubesnouts combined with SOs, still others have chin swellings resembling truncated SOs, while blunt nose species lack any facial protrusion ^41^. An obvious question raised by the present study is whether these diverse craniofacial morphologies bear any systematic relationships to the size and regional differentiation of the cerebellum.

## Methods

### Experimental Model and Subject Details

Adult fish (11-19 cm in length) of the species *Gnathonemus petersii* were used for experiments. Sex was not determined. Fish were housed in 60-gallon tanks in groups of 5-20. Water conductivity was maintained between 60-100 μS both in the fish’s home tanks and during experiments. All experiments performed in this study adhere to the American Physiological Society’s Guiding Principles in the Care and Use of Animals and were approved by the Institutional Animal Care and Use Committee of Columbia University.

### Surgery

Fish were anesthetized with MS-222 (1:25,000) and held against a foam pad. Aerated water containing MS-222 was continually passed over the fish’s gills for respiration throughout the surgery. The skin on the dorsal surface of the head was removed, and a long-lasting local anesthetic (0.75% Bupivacaine) was applied to the wound margins. A plastic rod was attached to the skull with Metabond (Parkell) to secure the head, and a craniotomy was performed over the C1 region of the cerebellum.

### Neuroanatomical Tracing

Following craniotomy surgery, and while the fish was still under anesthesia, a 10% solution of biotinylated dextran amine (BDA, Molecular Probes Europe, Leiden, The Netherlands) was iontophoretically delivered into the C1 region through a glass micropipette (inner tip diameter 20–30 μm) using a 5 μA positive pulsed direct current (7 s on/off). Subsequently, a 2% solution of Fluoro-Gold (FG, Fluorochrome, Englewood, CO) in 0.1 M cacodylate buffer (pH 7.3) was injected into the muscles of the Schnauzenorgan (SO). After a survival period of 5-7 days, the fish were anesthetized with MS-222 and transcardially perfused with 0.1 M PBS followed by 4% paraformaldehyde.

### C1 Microstimulation

Microstimulation was performed in head-fixed fish held lightly against a foam pad and sedated with Metomidate hydrochloride (Aquacalm) added to the tank water and passed continuously through the gills with a peristaltic pump (3.5 mg/l). Under these conditions fish continued to emit EODs at a low regular rate (∼1 Hz) ^42^. Microstimulation was performed with low-impedance tungsten or platinum-iridium microelectrodes (MicroProbes) using brief (0.1 ms duration) high-frequency pulse trains (typically 10 pulses at 200 Hz). Thresholds for SO movement in C1 were ∼5 μA. Sites were considered unresponsive if no movements were evoked by a 50 μA stimulus. Consistent with prior studies of cerebellar microstimulation ^43,44^, stimulation thresholds were lowest in the white matter just below the Purkinje cell layer. Responses to at least 5 consecutive stimulus repetitions were obtained for each site or parameter combination. SO movements were recorded with a camera positioned in front of the fish, recording at 166 or 300 fps using a USB 3.0 FLIR Chameleon Camera, with two mirrors providing side and bottom views. Stimulation triggered an LED visible to the camera for synchronization controlled by Bonsai (v2.8.2). Seven keypoints along the SO were tracked using DeepLabCut (v2.3rc1) for analysis of SO movement kinematics ^45^. Kinematic parameters of SO movement, including direction, amplitude, speed, and curvature, were extracted from video recordings. Movements of other body parts (including the mouth, tail, trunk, and pectoral fins) were also documented and, in a subset of experiments, recorded with a second camera. Stimulation sites were reconstructed histologically from serial sections (50-60 μm) based on a subset of penetrations in which the electrode was coated with the fluorescent dye DiI.

### Electrophysiological recordings

Following surgery, Gallamine triethiodide (Flaxedil; ∼20 μg/cm of body length) was administered to immobilize the fish, and fresh aerated water was continually passed over the fish’s gills for respiration. The rate of the electric organ discharge (EOD) motor command was monitored continuously by electrodes positioned near the electric organ in the tail. High-density multi-site extracellular recordings were performed using the Neuropixels Ultra probe ^24^. Probes were coated with fluorescent dye DiI (Invitrogen) and slowly inserted (20 μm/s) into the C1 region (∼3500 μm below the brain surface). Recordings were conducted using the acquisition software SpikeGLX (v20240129-phase30) in tip reference mode. The recorded data were processed to align channel sampling using the common average referencing software CatGT. Putative neuronal units were spike-sorted using Kilosort 2.0 and Phy2, and manually curated in Phy2. Passive displacement of the SO was performed using a motorized micromanipulator (Sensapex, uMP-4) coupled to the SO via plastic tubing. Electrosensory stimuli were delivered via a dipole electrode positioned near the SO or mouth using two sets of stimulation parameters designed to preferentially activate either low-frequency ampullary receptors mediating passive electrosensation (5 μA, 100 ms duration pulses delivered at 4 Hz) or high-frequency mormyromast receptors mediating active electrolocation (50 μA, 300 μs duration delivered time-locked to the fish’s EOD motor command).

### Neural data analysis

Neural responses to SO displacements (a sequence of eight movement direction repeated 12 times) were quantified using a modified version of the cortex-lab/spikes toolbox (https://github.com/cortex-lab/spikes). Briefly, spike counts were binned using a 10 ms moving window and then smoothed with a Gaussian filter. The filter was normalized to ensure its coefficients summed to one, preserving the overall signal amplitude. In order to cluster units based on their responses to SO displacement a recurrent neural Network (RNN) autoencoder was employed to reduce the dimensionality and extract features of the neuronal responses. K-means clustering was then applied to these features to identify patterns within the neural activity. The optimal number of clusters was determined using the silhouette method, which evaluates clustering quality by measuring how similar an object is to its own cluster compared to other clusters. Finally, a supervised RNN classifier was trained using the cluster assignments as pseudo-labels, allowing the prediction of cluster membership based on input sequences.

A Generalized Linear Model (GLM) was applied to predict SO position in two dimensions based on the neural activity of various subsets of recorded C1 units using a MATLAB subroutine (R2021b). The GLM was trained using data from 11 trials, and its performance was evaluated on the remaining trial. This leave-one-trial-out cross-validation process was repeated for each trial, generating 12 predictions in total.

### Behavioral characterization of C1 lesions

Foraging was filmed from the side in the fish’s home tank in a 15 × 30 × 12 cm arena equipped with top and front mirrors to provide additional views. The arena had three entrances allowing the fish to enter and exit at will. Experiments were performed during the light cycle and recorded at 300 fps using a USB 3.0 FLIR Chameleon Camera. Foraging was studied under two conditions: “floor” where fish searched for a single live blackworm placed on the arena floor and “trough” where a single blackworm was located inside a rectangular plastic trough (1 × 1 × 1.3 cm), requiring the fish to use its SO to extract the prey. Video recordings were begun after fish were acclimatized to the setup and efficiently captured worms under both conditions. At least five once-daily sessions consisting of at least 16 trials each were collected before and after lesions. In some cases, fish were followed for up to 2 weeks post-lesion. Lesions were made with low-impedance platinum-iridium microelectrodes (MicroProbes) using DC current (−50 μA for ∼100 seconds) at sites where SO movements could be evoked with low threshold current (<10 μA). All lesion sites were verified to be restricted to C1 by inspection of sagittal sections cut at 50-60 microns and counterstained with neutral red. The procedure for sham lesions was identical except that the stimulus isolator was switched off. For three fish included in the sham group we intended to place a lesion in the C3 region of the cerebellum (dorsal to C1) but post-hoc histology showed the lesion procedure failed completely.

DeepPoseKit (v0.3.6) in conjunction with custom bundle adjustment algorithms was used for 3D reconstruction of fish pose during foraging ^46,47^. Our custom bundle adjustment algorithm, inspired by Anipose (Anipose: A toolkit for robust markerless 3D pose estimation) ^48^, involved informed initialization of extrinsic and intrinsic camera parameters, to achieve reliable 3D coordinates. We filtered out low-confidence keypoints and performed temporal interpolation in 3D space to “infill” their position. A training dataset was generated consisting of over 500 frames consisting of seventeen labeled body parts, including five points along the length of the SO, for each camera view (for a total of 51 keypoints per fish) (Video 1). Fish kinematics were derived from the 3D coordinates of the fish’s body. Body orientation was determined frame-by-frame using the anterior head and middle of the body as reference points. Thrust, slip, and elevation components were calculated as the difference in head-middle position between successive frames along the axes of this reference frame. Roll was determined using the pectoral fin axis. The yaw and pitch components of the SO and yaw components of head and tail movements were calculated as the angle between orientations in a frame defined by the three key points describing the component (e.g., SO tip, mouth, and head for SO angles). SO and tail curvatures were calculated based on sets of five or three 3D points, respectively, each representing a location along the structure. From these, the cumulative arc length, radius of curvature, and curvature vector for each point were computed. Velocities were calculated based on the distances or angles traveled between two successive video frames.

### Quantification and Statistical Analysis

Data were analyzed off-line using custom Python (3.7) and Matlab code (Mathworks, Natick, MA). Non-parametric tests were used for testing statistical significance. Differences were considered significant at P < 0.05.

## Acknowledgements

This work was supported by grants from the NSF (IOS 2115007) and the NIH (NS075023) to N.B.S. We thank LF Abbott, Salomon Muller, Ashok Litwin Kumar, Liam Paninski, John Cunningham and members of the Sawtell laboratory for valuable discussions and comments on the manuscript.

## Author Contributions

F.P., M.G., D.N., P.C., D.E.E., and N.B.S. performed the experiments. F.P., M.G., D.N., P.C., D.E.E., D.B. and N.B.S. analyzed data and edited the manuscript. N.B.S. designed the project and wrote the manuscript.

## Supplementary Materials

**Figure S1:**
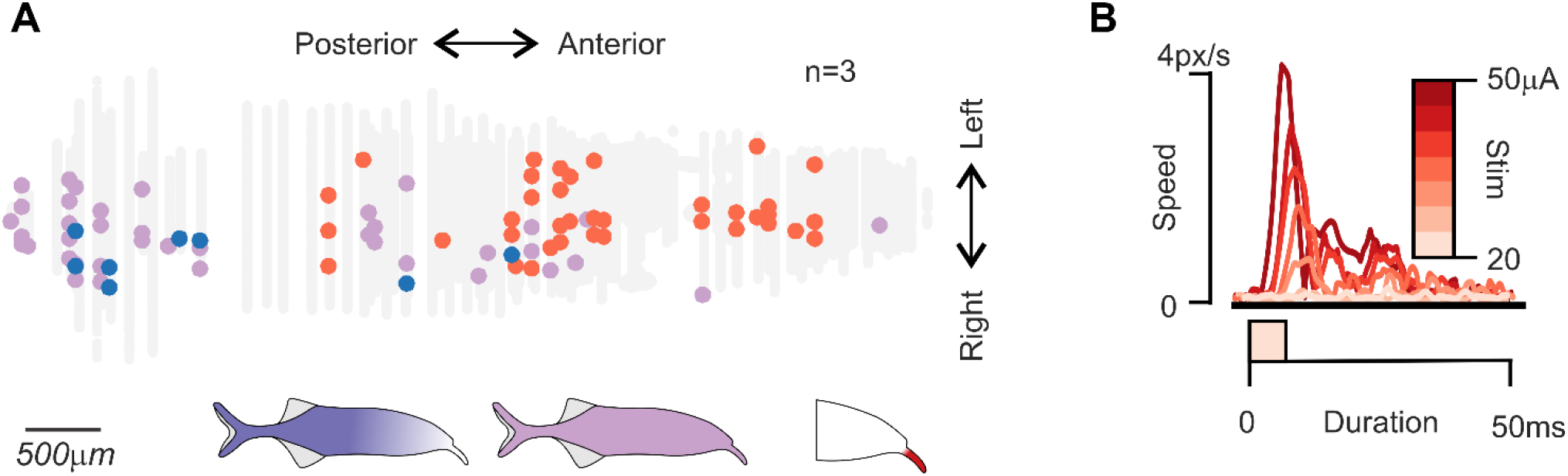
(A) Flatmap representation of the C1 region with color-coded circles indicating sites at which threshold stimulation evoked SO movements only (red), tail movements only (blue), or movements of both the SO and other body parts including the tail, trunk, or fins (purple) (n = 3 fish, 83 sites). (B) Effects of grading stimulation current strength on SO movement speed at a single site.

**Figure S2:**
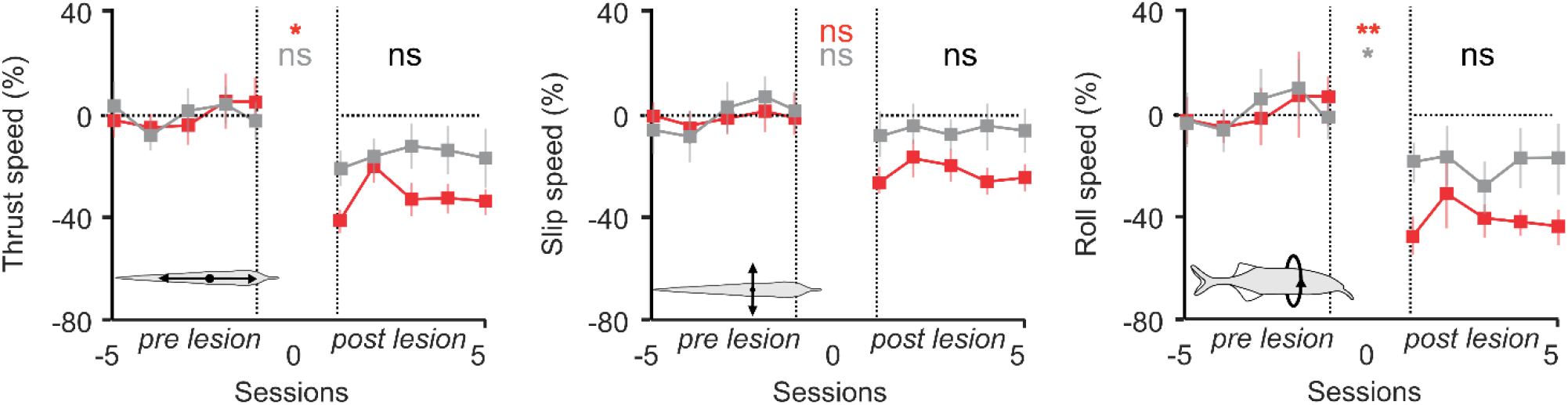
Thrust, Slip and Roll speed before and after C1 lesions in the *trough* condition for C1 lesioned (*n* = 10) and sham-lesioned fish (*n* = 8). Thrust speed: post C1 lesion vs post sham-lesion: Mann-Whitney test, p=0.90; pre vs post C1 lesion: Wilcoxon test, p=0.02; pre vs post sham lesion: Wilcoxon test, p=0.05. Slip speed: post C1 lesion vs post sham-lesion: Mann-Whitney, p=0.11; pre vs post C1 lesion: Wilcoxon test, p=0.20; pre vs post sham lesion: Wilcoxon, p=0.11. Roll speed: post C1 lesion vs post sham-lesion: Mann-Whitney, p=0.35; pre vs post C1 lesion: Wilcoxon test, p=0.005; pre vs post sham lesion: Wilcoxon, p=0.015.

**Video S1:** Left, video illustrating the match between the 2D tracked points (crosses) and the re- projection of the reconstructed 3D body parts (circles). Right, the inferred fish and camera positions after Bundle Adjustment match the geometry of the setup and the momentary fish pose, as seen in the left panel.

**Video S2:** Foraging under *floor* conditions before and after a unilateral C1 lesion. Note, the post-lesion reduction in SO movement and the shift in SO position contralateral to the side of the lesion.

**Video S3:** Foraging under *trough* conditions before and after a unilateral C1 lesion. Note, the increased time required to extract the worm from the trough post-lesion.

